# Spatial transcriptomics profiling of gallbladder adenocarcinoma: a detailed two-case study of progression from precursor lesions to cancer

**DOI:** 10.1101/2024.02.27.582232

**Authors:** Sophie Pirenne, Fátima Manzano-Núñez, Axelle Loriot, Sabine Cordi, Lieven Desmet, Selda Aydin, Catherine Hubert, Sébastien Toffoli, Nisha Limaye, Christine Sempoux, Mina Komuta, Laurent Gatto, Frédéric P. Lemaigre

## Abstract

**Background:** Most studies on tumour progression from precursor lesion toward gallbladder adenocarcinoma investigate lesions sampled from distinct patients, providing an overarching view of pathogenic cascades. Whether this reflects the tumourigenic process in individual patients remains insufficiently explored. Genomic and epigenomic studies suggest that a subset of gallbladder cancers originate from biliary intraepithelial neoplasia (BilIN) precursor lesions, whereas others form independently from BilINs. Spatial transcriptomic data supporting these conclusions are missing. Moreover, multiple areas with precursor or adenocarcinoma lesions can be detected within the same pathological sample. Yet, knowledge about intra-patient variability of such lesions is lacking.

**Methods:** To characterise the spatial transcriptomics of gallbladder cancer tumourigenesis in individual patients, we selected two patients with distinct cancer aetiology and whose samples simultaneously displayed multiple areas of normal epithelium, BilINs and adenocarcinoma. Using GeoMx digital spatial profiling, we characterised the whole transcriptome of a high number of regions of interest (ROIs) per sample in the two patients (24 and 32 ROIs respectively), with each ROI covering approximately 200 cells of normal epithelium, low-grade BilIN, high-grade BilIN or adenocarcinoma. Human gallbladder organoids and cell-ine derived tumours were used to investigate the tumour-promoting role of genes.

**Results:** Spatial transcriptomics revealed that each type of lesion displayed limited intra-patient transcriptomic variability. Our data further suggest that adenocarcinoma derived from high-grade BilIN in one patient and from low-grade BilIN in the other patient, with co-existing high-grade BilIN evolving via a distinct process in the latter case. The two patients displayed distinct sequences of signalling pathway activation during tumour progression, but Semaphorin 4A (*SEMA4A*) expression was repressed in both patients. Using human gallbladder-derived organoids and cell line-derived tumours, we provide evidence that repression of *SEMA4A* promotes pseudostratification of the epithelium and enhances cell migration and survival.

**Conclusion:** Gallbladder adenocarcinoma can develop according to patient-specific processes, and limited intra-patient variability of precursor and cancer lesions was noticed. Our data suggest that repression of *SEMA4A* can promote tumour progression. They also highlight the need to gain gene expression data in addition to histological information to avoid understimating the risk of low-grade preneoplastic lesions.

## Background

Gallbladder cancer accounts for less than 2% of cancer-related deaths and is often fortuitously diagnosed in gallbladder samples following cholecystectomy. The prognosis of the disease remains poor because patients often present at an advanced stage with unresectable tumour. Late diagnosis results from the lack of specific symptoms and of screening strategies, as well as from limited knowledge of the mechanisms driving tumour progression [1, 2]. Several studies investigated the pathology, genomics and epigenomics of tumour progression from precursor to cancer stage. They mostly investigated precursor and cancer lesions from distinct patients, precluding a good understanding of tumour progression at the individual level. Spatial transcriptomic data on precursor and adenocarcinoma lesions coexisting in a same patient are expected to provide clues on the mechanisms of tumour progression.

Adenocarcinomas account for >90% of gallbladder cancers and are considered to develop according to a metaplasia-dysplasia-adenocarcinoma histogenic sequence, in which the dysplastic stage consists of low-grade and high-grade biliary intraepithelial neoplasia (BilIN) [3–8]. BilINs consist of microscopic, flat or micropapillary lesions whose grade depends on the highest degree of cytological and architectural atypia. Low-grade BilINs display moderate cytoarchitectural atypia with pseudostratification of the nuclei, increased nucleo-cytoplasmic ratio and hyperchromasia. High-grade BilINs, formerly called carcinomas *in situ*, are defined by loss of nuclear polarity, marked cytological atypia and complex architectural patterns such as micropapillae [9–11].

Genomic alterations are already found at the BilIN stage. *KRAS* and *TP53* mutations were found in BilINs [12, 13] and a progressive increase in TP53 overexpression was proposed to occur during the evolution from low-grade BilIN to GBC [14]. A recent exome sequencing study uncovered *CTNNB1*, *TP53*, *ARID2* and *ERBB3* as the most frequently mutated genes in low-grade and high-grade BilINs [15]. When the disease evolves to invasive adenocarcinoma, alterations accumulate, and tumours display significant cell-type heterogeneity [16, 17]. At that stage the most frequent mutations affect *KRAS, CTNNB1, TP53*, *PI3KCA*, *ERBB2*, *CDKN2A* and *CDKN2B* [18–26], indicating that a fraction of the mutations found at the cancer stage can be detected in BilIN lesions. At the epigenome level, cancer lesions were split in subtypes with distinct hypermethylation:hypomethylation ratios; progressive and cumulative changes in promoter methylation were detected during progression from cholecystitis to cancer [26–29]. Increased hypermethylation was observed in adenocarcinomas as compared with BilINs. These epigenomic changes impacted Wnt/β-catenin signalling, Hedgehog signalling, tumour suppression and cell-microenvironment interactions [30–32]. Further, since gallstone-induced chronic inflammation drives gallbladder carcinogenesis [33], several authors compared the transcriptome of normal gallbladder tissue, gallbladder cancer, and gallbladder tissue exposed for varying lengths of time to gallstones, and identified molecular signatures associated with disease progression [34, 35]. Finally, in line with the genomic and epigenomic studies, single gene analyses revealed aberrant expression levels of TP53, P21, cyclin D1, EZH2, SMAD4 and CDKN2A protein at the BilIN stage [11], as well as the ability of a combined activation of KRAS and canonical Wnt/β-catenin or Notch signalling to induce gallbladder BilINs with malignant potential [36, 37]. Spatial transcriptomic data investigating BilIN to adenocarcinoma progression are still lacking.

Considering the genomics of tumour progression, Lin and coworkers provided evidence for patient-specific tumourigenic processes [15]. Their results indicated that precursor and cancer lesions within the same patient bear similar mutations, whereas the mutational signatures significantly vary between patients. Phylogenetic analysis of single nucleotide variants in lesions generated revealed that gallbladder cancer developed either BilIN-dependently or BilIN-independently [15].

To address the spatial transcriptomics of gallbladder tumour progression in individual patients, we selected samples from two patients displaying simultaneously several areas of gallbladder BilINs and adenocarcinoma, and collected an extensive spatial transcriptomic data set of each type of lesion per patient. The two patients were selected because of their differing cancer aetiology, offering the possibility to address intra-patient variability and tumour progression in distinct contexts. Our results show that each type of lesion displayed limited variability within the same patient, but significantly differed among patients. This revealed that the two patients have distinct tumourigenic processes, thereby corroborating earlier conclusions at the transcriptomic level. Our molecular investigations using gallbladder organoids also provide evidence that Semaphorin 4A (*SEMA4A*) repression, which was observed in the two patients, can contribute to tumour progression.

## Methods

### Spatial transcriptomics

Spatial profiling was performed on formalin-fixed paraffin-embedded (FFPE) tissue sections using GeoMx (NanoString Technologies, Seattle, WA, USA) [38] which was implemented by NanoString. The GeoMx Whole Transcriptome Atlas assay probe cocktail containing 18,677 probes was tested. Regions of interest (ROIs) subjected to spatial transcriptomic profiling encompassed epithelial areas of approximately 200 cells. The 24 ROIs of Patient #1 were all located on the same tissue section. For Patient #2, the ROIs were partitioned over two sections, namely 8 ROIs covering normal epithelium on one section, and 24 ROIs covering lesional tissue on a second section (Table 1). Additional information is provided in Supplementary Material (Supplementary methods).

**Table 1.**
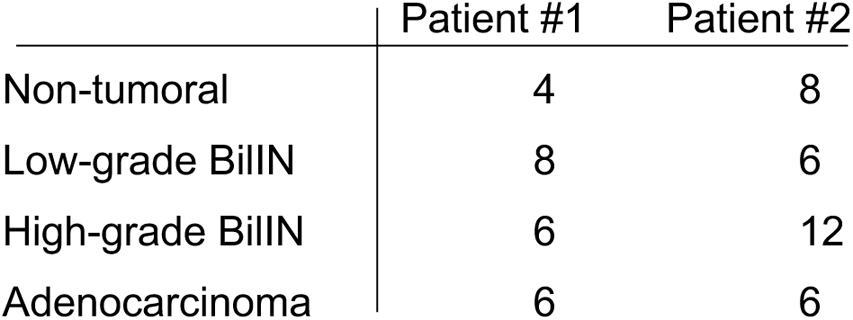
Number of ROIs subjected to spatial transcriptomic analysis.

|  | Patient #1 | Patient #2 |
| --- | --- | --- |
| Non-tumoral | 4 | 8 |
| Low-grade BillN | 8 | 6 |
| High-grade BillN | 6 | 12 |
| Adenocarcinoma | 6 | 6 |

#### Histology and staining

Hematoxylin/Eosin (H&E) staining was performed on 6 µm sections of FFPE tissues or organoids. Briefly, tissue sections were deparaffinised 3x 3 min in xylene, 3 min in 99%, 95%, 70% and 30% ethanol and deionised H_2_O. The sections were stained 7 sec in 100% hematoxylin, rinsed with H_2_O, stained for 7 sec in 100% eosine, and rinsed with deionised H_2_O. Dehydration of sections was performed in deionised H_2_O, followed by 30%, 70%, 95%, 99% ethanol for 30 sec, and 30 sec in xylene. Coverslips were placed on slides using Depex mounting medium (VWR, Leuven, Belgium). Pictures were taken with panoramic P250 Digital Slide Scanner (Histogenex, Antwerpen, Belgium) using 3DHISTECH Case Viewer software.

#### Immunofluorescence and immunohistochemistry

Immunofluorescence and immunohistochemistry were performed on 6 μm sections of FFPE tissues. FFPE tissue sections were deparaffinised 3x 3 min in xylene, 2 min in 99%, 95%, 70% and 30% ethanol and deionised water. Antigen retrieval was performed by the use of Lab Vision PT Module (Thermo Fisher Scientific, Waltham, MA), in 10 mM citrate pH 6. Sections were permeabilised for 10 min in 0.3% Triton X-100 in PBS before blocking for 1 h in 5% HS,10% BSA, 0.3% Triton X-100 in PBS. Primary antibodies were diluted in blocking solution at 4°C overnight and secondary antibodies were diluted in 10% BSA, 0.3% Triton X-100 in PBS at 37°C for 1 h. Images were taken with panoramic P250 Digital Slide Scanner (Histogenex, Antwerpen, Belgium) using 3DHISTECH Case Viewer software. Primary and secondary antibodies are described in Supplementary Material (Supplementary Table S1).

#### RNAscope *in situ* hybridisation

RNAScope RNA *in situ* hybridisation was performed on 5 μm sections of FFPE tissues, according to the manufacturer’s protocol for manual RNAscope®2.5 HD Assay—RED (#322360, Advanced Cell Diagnostics/Bio-Techne, Abingdon, United Kingdom). The tissue sections were incubated at 60°C for 1h30, deparaffinised 2x 5 min in xylene and dehydrated 2x 2 min in 99% ethanol. Endogenous peroxidase was blocked with hydrogen peroxide for 10 min at room temperature followed by two short washings with deionised water. Slides were heated for 10 sec at 100°C in deionised water, and antigen retrieval was performed for 15 min at 100°C using RNAscope®Target retrieval. Tissue sections were washed in deionised water and 99% ethanol. Slides were dried for 5 min at room temperature and tissues were delineated using an ImmEdge Hydrophobic Barrier Pen (#310018, Advanced Cell Diagnostics/Bio-Techne, Abingdon, United Kingdom). Slides were incubated for 15 min with RNAscope®Protease plus (diluted at 1/5 in deionised water) at 40°C, washed with deionised water and incubated with the Hs-COL1A1-Homo sapiens collagen type I alpha 1 mRNA probe for 2 h at 40°C. The tissue sections were washed with RNAscope®Wash buffer and six amplifications were performed (using six reagents AMP1-AMP6). The signal detection followed using RNAscope®Fast A and B reagents for 10 min at RT. The slides were kept in phosphate-buffered saline (PBS) overnight and immunostaining was performed: sections were blocked for 45 min at room temperature in 3% milk, 10% bovine serum albumin (BSA), 0.3% Triton in PBS. Primary and secondary antibodies were diluted in 10% BSA, 0.3% Triton in PBS. Primary antibodies were incubated overnight at 4°C and secondary antibodies were incubated 1h30 at 37°C. Pictures were taken with Cell Observer Spinning Disk (Carl Zeiss, Zaventem, Belgium) and analysed with Zen blue software. Primary and secondary antibodies are described in Supplementary Material (Supplementary Table S1).

### Gallbladder organoid culture

Human non-tumoral gallbladder tissues were obtained from patients who underwent cholecystectomy at the Cliniques Universitaires Saint-Luc, Brussels, using the method of Rimland and coworkers [39]. The karyotype of the selected organoid line was normal and whole exome sequencing detected an ERBB3^R675G^ missense mutation at an allelic fraction of 0.021. To analyse the impact of blocking SEMA4A in gallbladder organoids, the latter were split and plated. After 24 h, SEMA4A antibody (IgG-SEMA4A, #14-1002-82 eBioscience/Thermo Fisher scientific, Brussels, Belgium) was added into the medium (10 μg/ml) and organoids were grown for 3 days. Additional information is provided in Supplementary Material (Supplementary Methods).

#### Bioinformatic analysis of spatial transcriptomic profiling data

Sequencing quality was assessed for each ROI. Raw number of reads ranged from 1750000 to 21875463. Alignments rates, sequencing saturation and RTSQ30 were respectively higher than 80%, 70%, and 98% in all ROIs. The percent of detected genes (*i.e.* genes with an expression value higher than the LOQ value, defined as the negative probes geometric means + 2 standard deviations) was evaluated per segment, to identify low-performing AOIs that should be removed. All ROIs were kept, as values ranged from 13.6% to 51.4%. Raw count normalisation and differential expression analyses were performed using DESeq2 Bioconductor package v1.32.0 [40]. The generalised linear model was fitted using the following design: type of lesion * patient. The lists of differentially expressed genes generated by DESeq2 were ranked on the t-statistic values, and KEGG and HALLMARK gene set enrichment analyses were performed using clusterProfiler v4.0.5 [41].

## RESULTS

### Selection of normal epithelium, BilIN and adenocarcinoma in samples of human gallbladder

Our goal is to characterise the spatial transcriptome of gallbladder lesions during progression from normal epithelium to adenocarcinoma. This required gallbladder samples that simultaneously contain non-tumoral (*i.e.* histologically normal) epithelium, low-grade BilIN, high-grade BilIN and adenocarcinoma, from patients with distinct cancer aetiology. Each lesion must be large enough to enable us to analyse the whole transcriptome of several regions of each type of lesion. Samples that met these critera from two patients were identified in the biobank of the Cliniques Universitaires Saint-Luc: Patient #1 was an 81 year old woman who underwent cholecystectomy to treat cholecystitis; adenocarcinoma was an incidental finding. Patient #2 was a 53 year old man affected with primary sclerosing cholangitis (PSC) whose gallbladder was resected following imaging that revealed a thickening of the gallbladder wall. Pathological diagnoses of non-tumoral epithelium, BilINs and adenocarcinoma were made on H&E-stained sections, and were confirmed by two expert pathologists. Patient #1 displayed two small foci of intestinal metaplasia, and no metaplasia was detected in Patient#2. GeoMx Digital Spatial Profiling (NanoString) [38] was implemented on sections adjacent to the H&E-stained sections to collect whole transcriptome data from 56 epithelial ROIs, each covering approximately 200 epithelial cells of non-tumoral epithelium, BilIN and adenocarcinoma (Table 1). Metaplasia in Patient #1 were too small for spatial profiling. Figure 1 illustrates the spatial distribution of areas in which ROIs were delineated (Fig. 1A), as well as examples of H&E-stained non-tumoral epithelium, BilINs and adenocarcinomas (Fig. 1B). Epithelial ROIs were delineated on sections stained with antibodies which detect markers of the epithelium (panCytokeratin), leukocytes (CD45), and mesenchymal cells (α smooth muscle actin). Nuclei were immunolabeled with anti-Human antigen R antibodies (Supplementary Fig. S1). The ROIs were subjected to transcriptomic analyses as described in Methods.

**Fig. 1.**
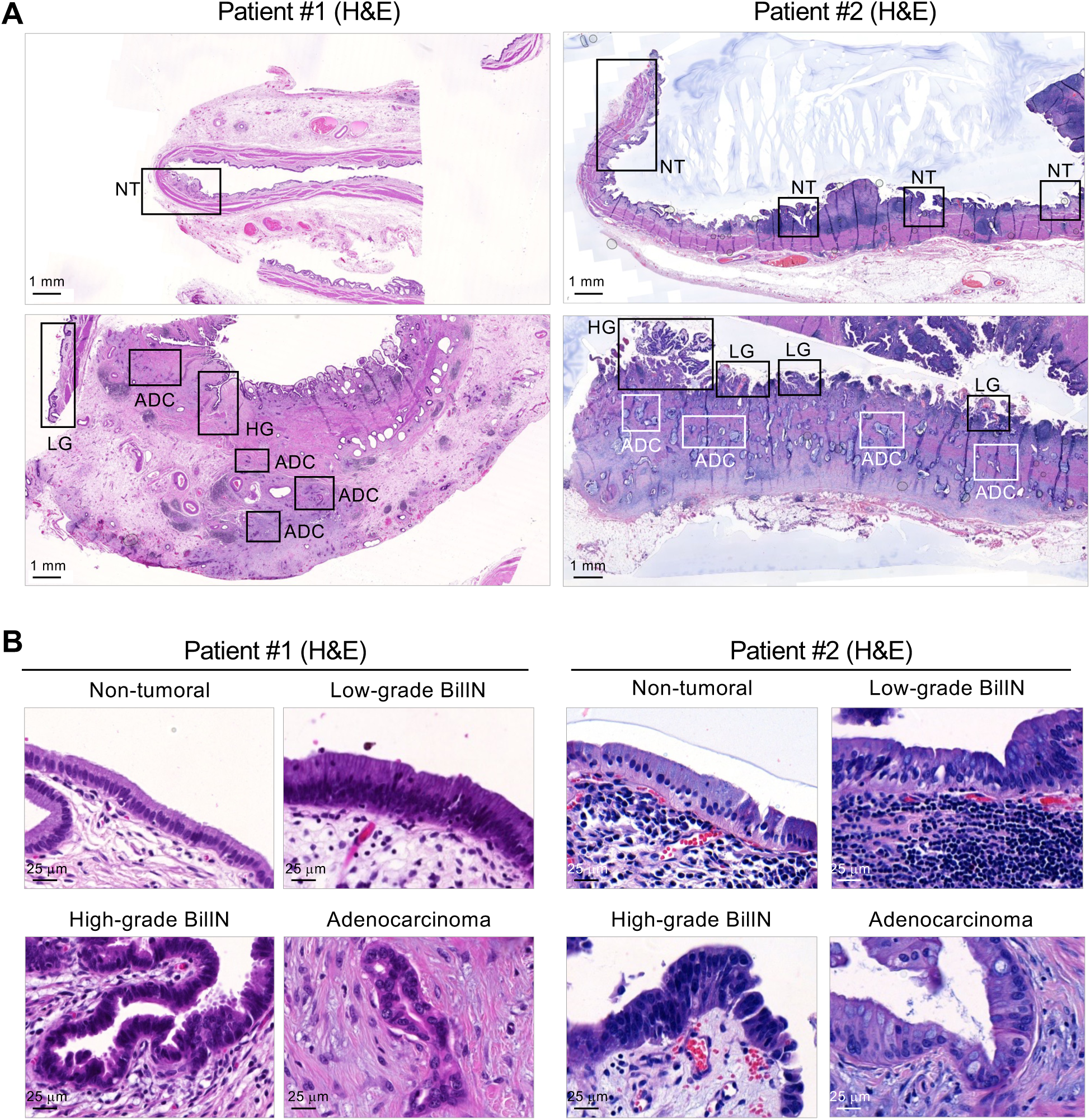
Selection of non-tumoral (histologically normal) epithelium, BilIN and adenocarcinoma in samples of human gallbladder. (A) Low magnification view of gallbladder sections. Squares indicate tissue areas in which several epithelial ROIs were delineated as shown in Supplementary Fig. S1. (B) Illustrative examples of non-tumoral epithelium, low-grade BilIN, high-grade BilIN and adenocarcinoma. ADC, area containing adenocarcinomas; H&E, haematoxylin-eosin; HG, area containing high-grade BilINs; LG, area containing low-grade BilINs; NT, area containing non-tumoral epithelium.

### Spatial transcriptomic analysis suggests limited intra-patient variability and distinct modes of tumour progression among the two patients

Principal component analysis (PCA) of the 56 transcriptomes revealed a remarkable clustering of the non-tumoral epithelial samples of the two patients (Fig. 2A). ROIs from the same type of lesions clustered together within the same patient, but were separated between patients. In Patient #1, adenocarcinoma ROIs clustered close to high-grade BilIN ROIs, whereas adenocarcinomas in Patient #2 appeared closely related to low-grade BilINs. These results were corroborated by the number of differentially expressed genes (log_2_ fold change ≥ 1.0; p_adj_ ≤ 0.05) when cross-comparing all tissue types (Supplementary Fig. S2A). Together, these data revealed that each lesional type displays limited intra-patient variability, but that distinct mechanisms are driving tumourigenesis in the two patients. Moreover, the PCA plot suggested that adenocarcinoma evolved according to a normal ® low-grade BilIN ® high-grade BilIN ® adenocarcinoma sequence in Patient #1, and according to a normal ® low-grade BilIN ® adenocarcinoma sequence in Patient #2, with high-grade BilIN emerging separately from adenocarcinoma in this patient.

**Fig. 2.**
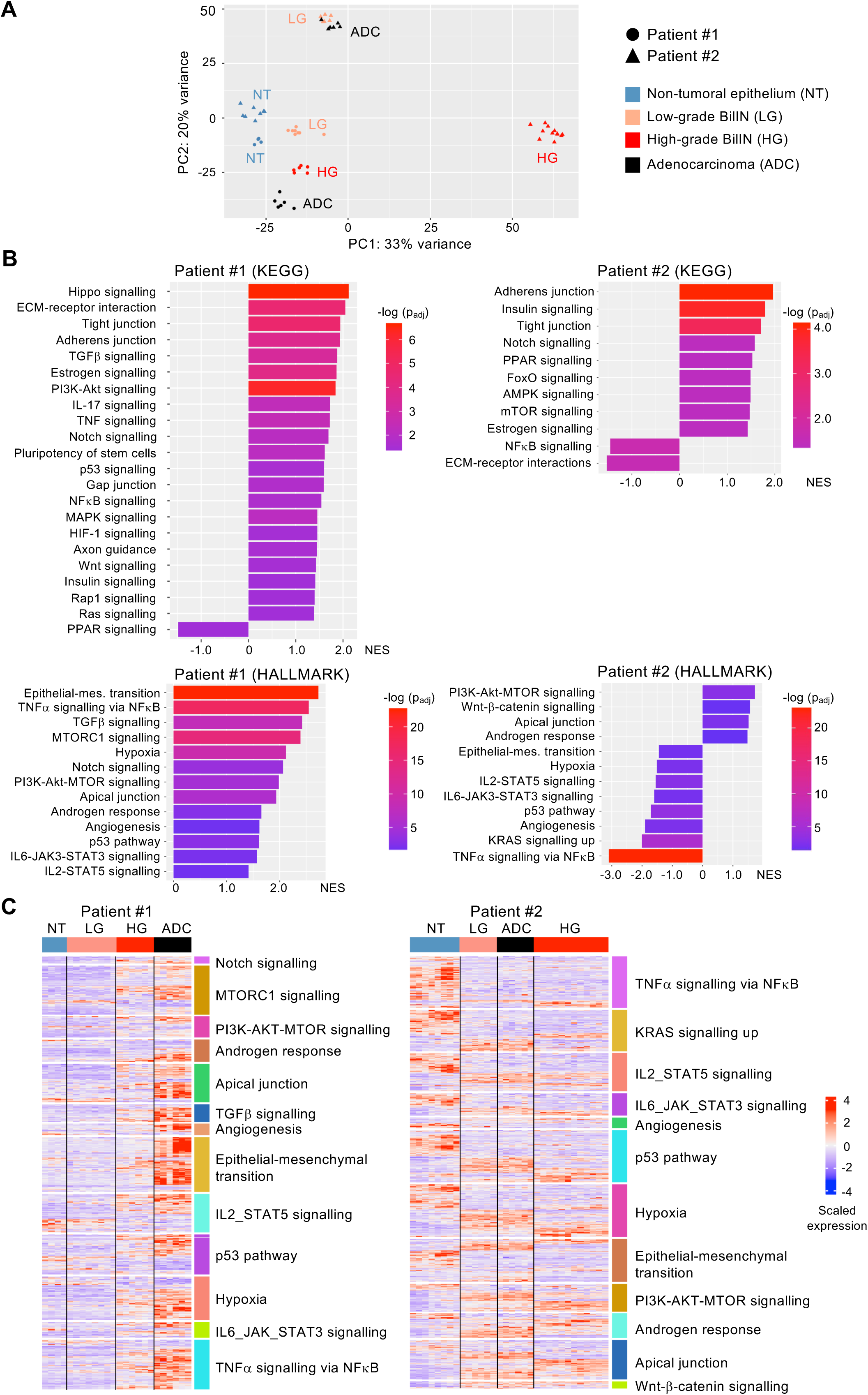
Distinct modes of tumour progression in two patients revealed by spatial transcriptomic analysis. (A) PCA plot of the whole transcriptome of 56 ROIs comprising non-tumoral (histologically normal) epithelia, low-grade biliary BilINs, high-grade BilINs and adenocarcinomas. (B) Heatmaps of GSEA enrichment scores using the KEGG pathway and HALLMARK gene sets (p_adj_ ≤ 0.05). (C) Heatmaps of genes from the HALLMARK gene sets that are differentially expressed between adenocarcinoma and normal epithelium ROIs (p_adj_ ≤ 0.05). ADC, adenocarcinoma; HG, high-grade BilIN; LG, low-grade BilIN; NES, normalised enrichment score; NT, non-tumoral epithelium.

We next compared the lesions in the two patients and focused on signalling pathways. Using Gene Set Enrichment Analysis (GSEA) [42], we found several enriched signalling pathways when comparing adenocarcinoma and non-tumoral epithelium. Negative or positive enrichment scores reflect enrichment of downregulated or upregulated genes, respectively (Fig. 2B). The use of KEGG or HALLMARKS gene sets revealed several pathways that were enriched in both patients, and other pathways that were enriched in only one patient. Heatmaps illustrate genes from the HALLMARKS and KEGG pathway gene sets that are differentially expressed between adenocarcinoma and non-tumoral epithelium in the two patients (Fig. 2C; Supplementary Fig.S2B).

Galbladder cancer is often associated with mutations in *PI3KCA, CTNNB1, KRAS, TP53*, and *ERBB2* [18–26]. GSEA revealed that PI3K-AKT-mTOR signalling (HALLMARK) is enriched in adenocarcinoma of both patients (Fig. 2C), and out of the 38 leading edge genes in Patient #2, 23 overlapped with the leading edge genes in Patient#1. HALLMARK gene sets are based on coordinately expressed and biologically relevant genes, and identify pathway activation phenotypes [43]. Therefore, the positive enrichment of PI3K-AKT-mTOR signalling reflects activation of the pathway. Further, GSEA suggested enrichment of Wnt signalling in both patients, when considering the KEGG Wnt signalling gene set in Patient #1 and the HALLMARK Wnt-β-catenin gene set in Patient #2 (Fig. 2C). However, the two gene sets differ in their composition, leading to different conclusions in the two patients. In Patient #1, Wnt ligands (*WNT7B*, *WNT8A, WNT10A*, *WNT11*), receptors (*FZD2*, *FZD5*) and effector (*TCF7L2*) were upregulated in adenocarcinoma as compared to non-tumoral epithelium. Genes induced by Wnt signalling and reflecting activation of a negative feedback loop (*AXIN2*, *GSK3B*) further reveal dynamic activity of the Wnt pathway in this patient (Supplementary Fig. S2C). In contrast, in Patient #2, only 13 genes from the HALLMARK Wnt-β-catenin gene set were significantly enriched. Among these, most genes are not typical for Wnt signalling and belong to pathways with which Wnt signalling crossreacts. *CTNNB1* is upregulated in adenocarcinoma of Patient #2 (log_2_ fold change=1.10; p_adj_=8.65x10^-10^), in parallel with upregulation of Wnt signalling inhibitors *DKK4* (log_2_ fold change=0.86; p_adj_=84.76x10^-4^) and *CSNK1E* (log_2_ fold change=0.51; p_adj_=1.66x10^-3^). Therefore, the analysis of genes of the HALLMARK Wnt-β-catenin gene set does not strongly support that Wnt signalling is active in Patient #2. KRAS signalling differs between the two patients, as evidenced by enrichment of RAS signalling (KEGG) in Patient #1, but downregulation of several KRAS targets within the KRAS signalling up gene set (HALLMARK) in Patient #2 (Fig. 2C). Similar to KRAS signalling, the p53 pathway differed between patients. Finally, GSEA did not highlight ERBB signalling. However, we found significant overexpression of EGFR, ERBB2 and ERBB3 in Patient #2, but only overexpression of ERBB2 in Patient #1 (Fig. 3A).

**Fig. 3.**
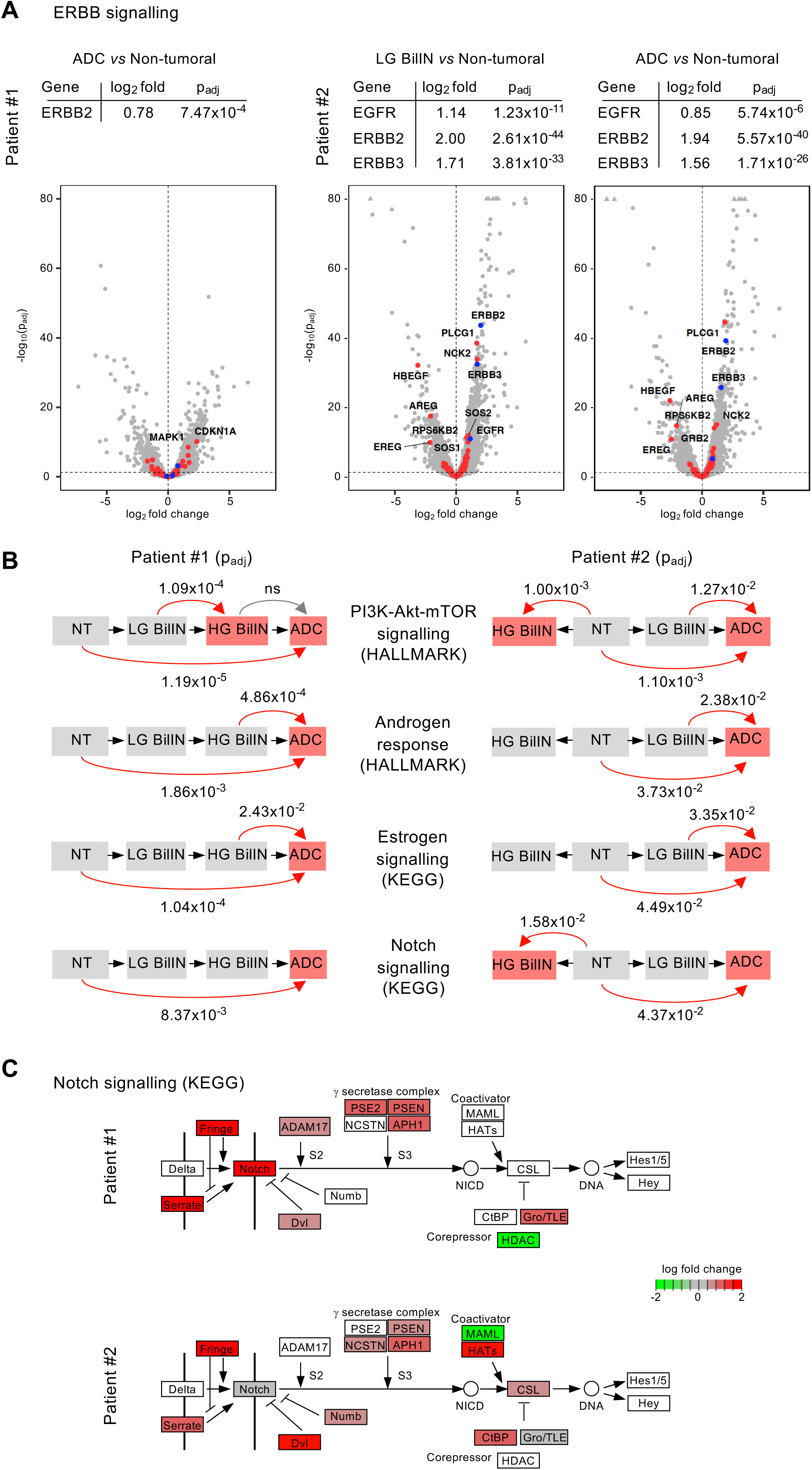
Distinct modes of signalling pathway activation in two patients revealed by spatial transcriptomic analysis. (A) Expression of ERBB receptors and ERBB signalling pathway genes during tumour progression. Tables mention the fold change inductions between lesions in the two patients. The corresponding volcano plots are shown, with blue dots highlighting EGFR/ERBB receptors. (B) Sequence of enrichment of signalling pathways during tumour progression as determined by GSEA using KEGG pathway and HALLMARK gene sets. Significant enrichments are indicated with p_adj_ values. Red boxes, lesions showing enrichment of the pathway. ns, not significant. (C) Differential expression of genes between adenocarcinoma and non-tumoral epithelium in the KEGG pathway Notch. ADC, adenocarcinoma; HG, high-grade BilIN; LG, low-grade BilIN; NES, normalised enrichment score; ns, non-specific; NT-non-tumoral epithelium.

Although both patients can display enrichment of the same pathway, we noticed that the sequence of enrichment during tumourigenesis may differ among the patients. Indeed, PI3K-AKT-mTOR signalling became enriched in precursor lesions of Patient #1, namely at the low-grade BilIN ® high-grade BilIN transition, whereas it became enriched only at the adenocarcinoma stage in Patient #2 (Fig. 3B). Other pathways whose enrichment is shared between the patients may in contrast display a similar sequence of enrichment. Indeed, androgen response and estrogen signalling became enriched at the precursor-to-adenocarcinoma transition (Fig. 3B). Notch signalling was also enriched in adenocarcinoma of both patients, and the enrichment was only significant when comparing non-tumoral epithelium and adenocarcinoma, not when comparing the precursor to adenocarcinoma transitions. This likely reflected a progressive activation throughout the tumourigenic process, without significant jumps between lesional states. Moreover, comparing the expression of leading edge genes in the Notch pathway also revealed interesting differences such as the strong upregulation of *NOTCH3* in Patient #1 (log_2_ fold change=2.05; padj= 5.2x10^-11^) and more modest upregulation of this gene in Patient #2 (log_2_ fold change=0.77; padj= 1.5x10^-3^) (Fig. 3C).

### Spatial transcriptomic analysis identifies hybrid epithelial and mesenchymal states

Nepal and coworkers considered the hallmark “epithelial-mesenchymal transition (EMT)” as indicative of poor prognosis [26]. In Patient #1, the corresponding HALLMARK gene set has the highest enrichment score when comparing adenocarcinoma with non-tumoral epithelium (Fig. 2B-C). The sequence of EMT enrichment is shown in Fig. 4A. No similar enrichment was found in Patient #2. Transcription factors typical for EMT and *CADHERINS* showed no significant differential expression during tumour progression in either patient (Fig. 4B). In contrast, extracellular matrix-coding genes were significantly upregulated (Fig. 4C). To support the latter data at the histological level, we resorted to RNAscope *in situ* hybridisation. We detected rare mRNAs coding for *COL1A1* in non-tumoral epithelia of the two patients. Strong induction of *COL1A1* was detected in high-grade BilIN of Patient #1, but also in low-grade BilINs of Patient #2 (Fig. 4D). We concluded that both patients displayed criteria of hybrid epithelial and mesenchymal states. The marked EMT in Patient #1 suggests that the adenocarcinoma belongs to the poor prognosis category.

**Fig. 4.**
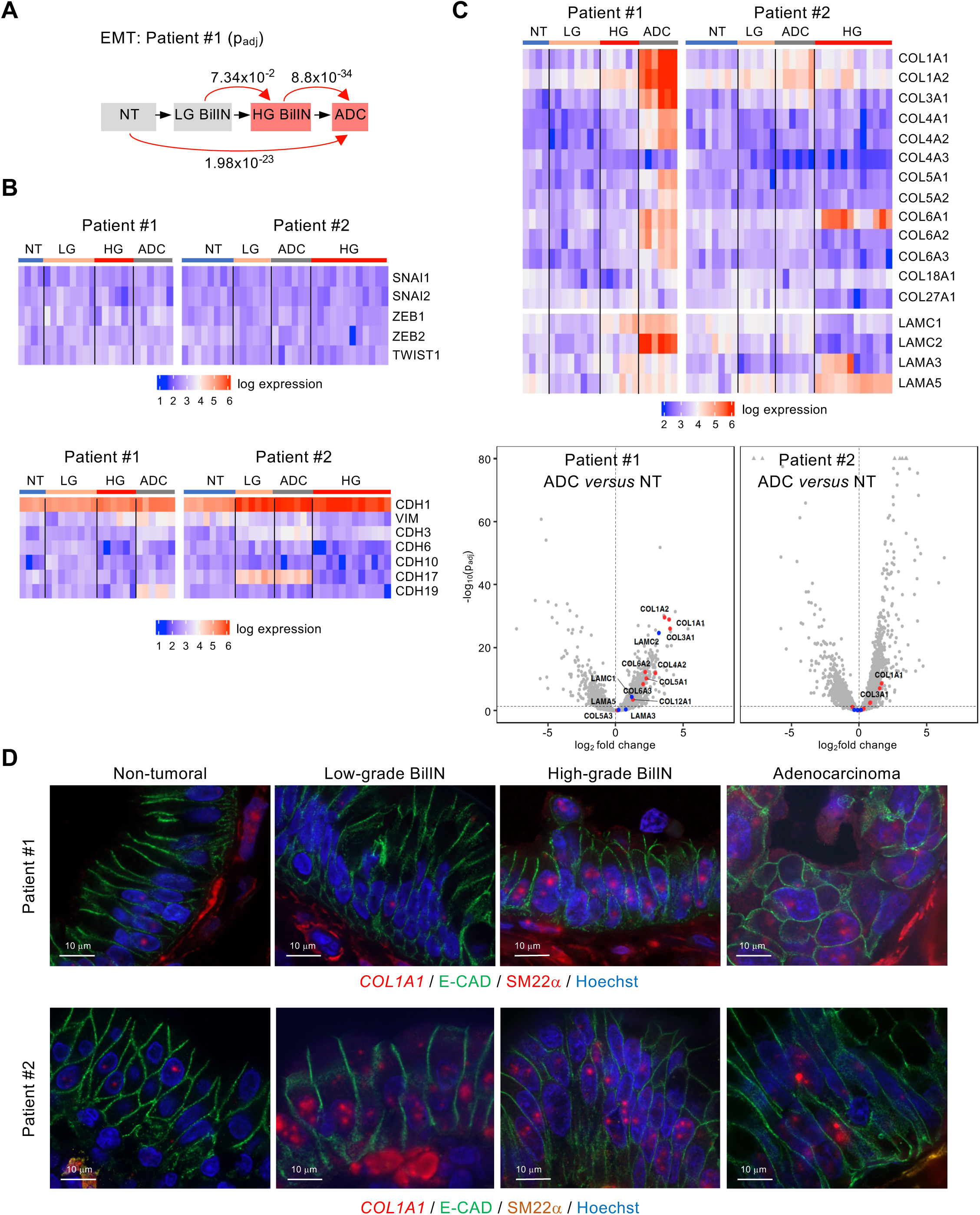
Hybrid epithelial-mesenchymal states during tumour progression. (A) Enrichment sequence of EMT (HALLMARK) in Patient #1 demonstrates enrichment throughout tumourigenesis. Significant enrichments are indicated with p_adj_ values. Red boxes, lesions showing enrichment of the pathway. (B) Gene expression heatmaps of EMT-promoting transcription factors, and of *VIMENTIN* and *CADHERINS* show little or no variation during tumourigenesis. (C) Heatmap and volcano plots showing *COLLAGEN* and *LAMININ* gene expression in the two patients. Blue dots in volcano plots indicate *LAMININ* genes. (D) RNAscope *in situ* hybridisation demonstrates induction of *COL1A1* mRNA (red dots) starting in high-grade BilINs in Patient #1 and in low-grade BilIN of Patient #2. Tissue sections were immunostained to mark epithelial cells (E-CADHERIN; E-CAD), nuclei (Hoechst), and mesenchymal cells (Smooth muscle protein 22α; SM22α). ADC, adenocarcinoma; HG, high-grade BilIN; LG, low-grade BilIN; NES, normalised enrichment score; NT, non-tumoral epithelium.

### SEMAPHORIN4A downregulation promotes tumour progression

Our GSEA data uncovered axon guidance signalling as a potential driver of tumour progression in Patient #1 (Fig. 2C). Axon guidance genes, including *SEMAPHORIN/PLEXIN* ligand-receptor pairs, were enriched in Patient #1 adenocarcinomas, but not in Patient #2 (Supplementary Fig. S3). *SEMA4A* was downregulated in the adenocarcinomas of both patients, and this was noticed already at the precursor stages (Fig. 5A). The involvement of SEMA4A in gallbladder cancer is unexplored, but *SEMA4A* loss-of-function mutation in familial colorectal cancer type X was found to promote cancer development, thereby revealing a tumour suppressor role for SEMA4A [44, 45]. This prompted us to investigate the role of SEMA4A in gallbladder cancer development.

**Fig. 5.**
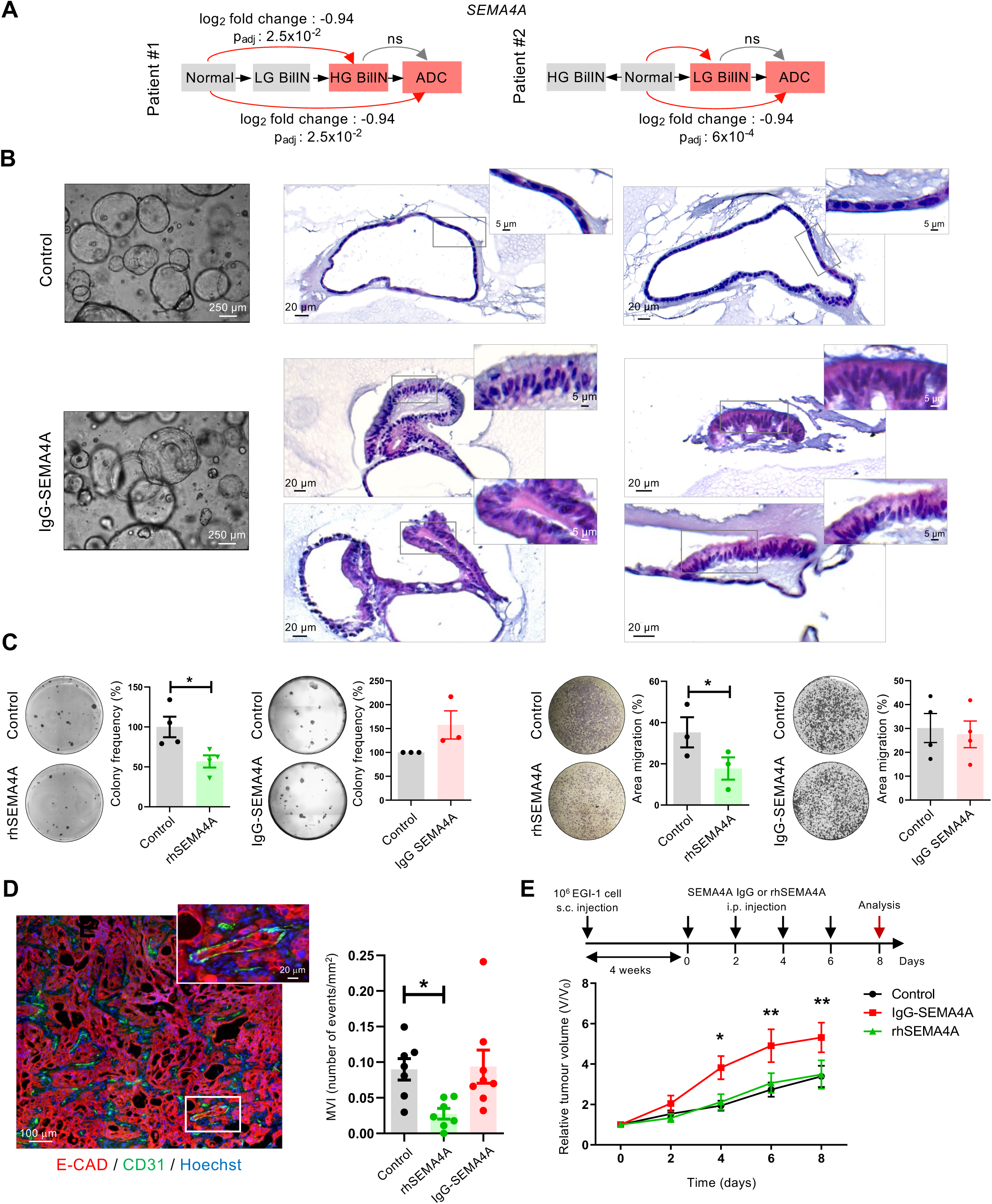
SEMA4A displays tumour suppressor properties. (A) *SEMA4A* gene expression is reduced during gallbladder cancer progression. ADC, adenocarcinoma; HG, high-grade BilIN; LG, low-grade BilIN; NES, normalised enrichment score; NT, non-tumoral epithelium; ns, non-specific. (B) The epithelium of gallbladder organoids treated with blocking anti-SEMA4A IgG antibody displays focal areas of pseudostratification. This effect was monitored in two experiments out of four. (C) rhSEMA4A reduces clonogenicity and transwell migration of cultured EGI-1 cells, whereas anti-SEMA4A IgG antibody had little or no effect. Data show means +/- SEM; n=3 or 4; statistical significance was calculated by applying a paired *t*-test (*, p<0.05). (D) Microvascular invasion (MVI) is illustrated in subcutaneous EGI-1 cell tumours following intraperitoneal injection of rhSEMA4A or of blocking IgG SEMA4A antibodies, according to the timing shown in panel E. The graph shows that rhSEMA4A reduces the number or MVI events in EGI-1 cell tumours. One-way ANOVA was used to compare means (*, p<0.05). For each condition, two mice were injected subcutaneously (4 tumours/mouse). (E) Growth of subcutaneous EGI-1 cell tumours following intraperitoneal injection of PBS (control), blocking anti-SEMA4A IgG antibody, or rhSEMA4A. n=10 (control), 8 (IgG SEMA4A) and 7 (rhSEMA4A). Relative tumour volume and SEM are plotted. Differences between groups were evaluated by performing a two-way Analysis of Variance (two-way repeated measures ANOVA) with Bonferroni correction (*, p<0.05; **, p<0.01). For further statistical validation, a random intercept-random slope model with continuous time was fitted. This showed a significant interaction between the time and group effect (p=0.03), in particular, the contrast between SEMA4A IgG and control is significant (p=0.048) but not that between control and rhSEMA4A (p=0.95).

We first generated organoids from gallbladder epithelium and selected a line which displayed no karyotypic anomalies. It expressed biliary-specific markers and exhibited biliary transport functions (Supplementary Fig. S4). It also expressed the genes coding for SEMA4A and its receptor Plexin B1 (PLXNB1) (Supplementary Fig. S3B). To mimick the downregulation of *SEMA4A* observed in our transcriptomic analyses, we incubated the organoids for 3 days with a blocking anti-SEMA4A IgG antibody. We found no change in cell proliferation, but observed local areas of pseudostratification of the epithelium in a subset of organoids (Fig. 5B). The histology of those areas was reminiscent of BilIN, indicating that inhibiting SEMA4A impacts cell polarisation.

We next determined if SEMA4A had additional tumour suppressor properties. Since the organoid lines were not able to induce tumour formation after subcutaneous injection in immunodeficient NSG mice, we used the human extrahepatic cholangiocarcinoma cell line EGI-1. *In vitro*, clonogenic and transwell migration assays demonstrated that adding rhSEMA4A to cultured EGI-1 cells reduced their clonogenicity and migration (Fig. 5C). Blocking anti-SEMA4A IgG antibody slightly but not significantly increased colony formation, and did not impact cell migration (Fig. 5C). *In vivo*, subcutaneous injection of EGI-1 cells in immunodeficient NSG mice resulted in the formation of tumours. Consistent with the decreased migration induced *in vitro* by rhSEMA4A, intraperitoneal administration of rhSEMA4A resulted in a significant reduction of microvascular invasion in EGI-1 cell-derived tumours (Fig. 5D). Anti-SEMA4A IgG antibody had no effect on microvascular invasion in the tumours. Recombinant SEMA4A did not impact tumour growth. In contrast, blocking IgG anti-SEMA4A antibody accelerated growth at the earliest stages of tumour growth to progressively reach a plateau (Fig. 5E). We conclude that SEMA4 can control tumour progression by impacting polarity, clonogenicity and migration of cells.

## DISCUSSION

Earlier mutational profiling of precursor and cancer lesions coexisting in a same patient provided evidence that adenocarcinoma development may be BilIN-dependent or - independent [15]. Here, using GeoMx technology we extended these findings at the transcriptional level in two patients. We showed that lesions exhibited low intra-patient variability, but exhibited patient-specific sequences of signalling pathway activation.

In Patient #1, ROIs from a same type of lesion were often located at a short distance from each other, except for adenocarcinoma ROIs which were more scattered throughout the tissue sample. In Patient #2, high-grade BilIN ROIs were close to each other, but low-grade BilIN, adenocarcinoma and non-tumoral epithelium ROIs were significantly dispersed (Fig. 1A). Still, in spite of the scattering within the tissue, the transcriptomic profile of lesions belonging to the same histological type showed low intra-patient variabilty. Such transcriptomic homogeneity likely reflects that cells from a same type of lesion proliferated in a similar environment and with limited accumulation of new mutations. Clonal analysis of gallbladder cancers revealed subclonal diversification [46], in line with significant epithelial cell heterogeneity in the adenocarcinoma lesions notices in single cell RNA sequencing studies [16, 17]. However, our patient samples contained all lesional types on the same tissue sections, suggesting that cancer lesions had not enough time to accumulate genomic lesions, invade the tissue and produce subclones.

The neighbourhood of low-grade BilIN, high-grade BilIN and adenocarcinoma which may occur in pathological samples, leads us to surmise that the epithelium undergoes a normal epithelium → low-grade BilIN → high-grade BilIN → adenocarcinoma histogenic sequence. *A contrario*, the transcriptomic profile of Patient #2 strongly suggests that adenocarcinoma derived from low-grade BilIN, not from adjacent high-grade BilINs. This contrasted with Patient #1 whose adenocarcinoma ROIs were closely related to high-grade BilINs. We excluded that adenocarcinoma in Patient #2 corresponded to low-grade BilINs extending in Rockitansky-Aschoff sinuses. In Patient #2, only 58 genes were 2-fold up- or downregulated when comparing low-grade BilIN and adenocarcinoma, revealing that low-grade BilIN may be at high risk for evolution towards invasive cancer.

Many signalling pathways were activated during tumour progression and several were common between the two patients. However, the sequence of pathway activation differed between patients, some of the common pathways being activated at the BilIN stage in one patient, but only in the adenocarcinoma cells in the other patient. Therefore, our work suggests that various combinations of pathway activations may end up yielding cancer, no specific pathway or combination of pathways being responsible for transition from one stage to the other.

The HALLMARK gene set “Inflammatory response” was enriched in adenocarcinomas of both patients (not shown), reflecting their common chronic inflammatory background. Still, the tumour aetiology differed in Patients #1 and #2, with Patient #2 being affected with PSC, a disease with high incidence of adenocarcinoma [47]. The adenocarcinoma in Patient#2 was mucosecreting (Fig. 1B), unlike the carcinoma in Patient #1. The mutational profile of cholangiocarcinoma in PSC is heterogeneous and affects genes similar to those in non-PSC associated cholangiocarcinoma, the most frequently mutated being *TP53*, *KRAS*, *PI3KCA* and *GNAS*. In low-grade and high-grade dysplastic lesions, loss or amplifications of several genes, as well as mutations in *ERBB2* and *TP53*, can already occur [48, 49]. Our work extend these data at the transcriptomic level and highlight that low-grade BilIN can be very closely related to adenocarcinoma.

EMT is a phenotypic continuum during which epithelial cells evolve to a mesenchymal state via transitional or hybrid states [50]. It involves disruption of polarity and intercellular adhesion, changes in the interaction between cells and extracellular matrix, and increased migration [51, 52]. Our RNAscope analysis of *COL1A1* expression demonstrated that signs of EMT are detectable early on in epithelial cells during tumour progression, reflecting the emergence of hybrid epithelial-mesenchymal states.

SEMA4A is a tumour suppressor in colorectal cancer [44, 45]. Here we found that it is downregulated in both patients during gallbladder tumour progression, starting at the BilIN stage. Gallbladder organoids expressed *SEMA4A* and its receptor PLXNB1 and the levels of *SEMA4A* expression varied considerably (Supplementary Fig. S3B), likely explaining the variable pseudostratification of the gallbladder organoids when treated with blocking IgG antibody (Fig. 5B). Also, the low levels of *SEMA4A* and *PLXNB1* in cholangiocarcinoma EGI-1 cells, as compared to organoids derived from normal gallbladder epithelium, fit with the notion that SEMA4A is repressed in biliary cancer cells and with our observation that anti-SEMA4 blocking antibodies have limited or no effect on clonogenicity and migration of EGI-1 cells *in vitro*. *In vivo*, we detected a higher level of *SEMA4A* in EGI-1 cell-drived tumours than in *in vitro* cultured EGI-1 cells (Supplementary Fig. S3B). We excluded that this results from SEMA4A production by tumour-invading mouse cells, as our PCR primers were designed to specifically detect human SEMA4A. Inhibiting this *in vivo* production of SEMA4A enabled us to monitor growth-promoting properties of anti-SEMA4 blocking antibodies. How these anti-SEMA4A antibodies promote EGI-1 cell-derived tumour growth remains unclear. Indeed, our data show that inhibiting SEMA4A accelerates tumour growth during 4 days. This effect slows down to reach a plateau (Fig. 5E), and at the plateau stage we noticed a slight but not significant increase in proliferation rate, as evidenced by immunostaining for phospho-Histone H3 (Supplementary Fig. S3C). We hypothesize that anti-SEMA4A antibodies promoted proliferation mainly during the first 4 days of treatment. Interestingly, rhSEMA4 did not impact tumour growth, but decreased microvascular invasion, suggesting that reduction of SEMA4 promotes metastasis. The signalling pathways mediating the effects of SEMA4A on migration, polarity and potentially proliferation deserve further investigation. Further studies will determine how frequently SEMA4A is repressed at early stages of gallblader cancer and whether understanding its pathway may lead to identify biomarkers of early diagnosis of gallbladder tumours.

## CONCLUSION

Our spatial transcriptomic analysis reveals that precursor and cancer lesions can display limited intra-patient variability during gallbladder cancer progression and supports that tumourigenic mechanisms are patient-specific. Repression of *SEMA4A* may contribute to tumour progression. Our work also underscores that low-grade BilINs may be at high risk for developing to cancer and should ideally be characterised by gene expression profiling.

## ABBREVIATIONS

BilIN: biliary intraepithelial neoplasia
EMT: epithelial-mesenchymal transition
FFPE: formalin-fixed paraffin-embedded
H&E: haematoxylin and eosin
PSC: primary sclerosing cholangitis
ROI: region of interest.

## DECLARATIONS

### Ethics approval

The study on human samples was conducted in compliance with the ethical guidelines of the 2013 Declaration of Helsinki and was approved by the Comité d’Ethique Hospitalo-Facultaire (UCLouvain and Cliniques Universitaires Saint-Luc) with numbers 2018/06Jul/281 and 2021/26OCT/444. In accordance with article 8 of the internal rules of the Cliniques universitaires Saint-Luc, the need for informed consent was waived to the present retrospective study. The study is based solely on the analysis of residual human body material and on the collection of data existing in the medical files of patients who have not expressed their opposition to the use of their medical file for scientific research purposes. An informed consent exemption request was thus presented to the Ethics Committee, which was accepted. Mice received humane care and the research protocol was approved by the Animal Welfare Committee of the Université Catholique de Louvain with number 2022/UCL/MD/17.

### Consent for publication

Not applicable.

### Data availability

Data are stored under Gene Expression Omnibus (GEO) accession number GSE259311

### Authors’ contribution

S.P., F.M.-N., A.L., M.K., L.G. and F.L. designed the study; S.P., F.M.-N., A.L., S.C., S.A., C.H., S.T., acquired data; S.P., F.M.-N., A.L., L.D., N.L., C.S., M.K., L.G. and F.L. analysed and interpreted data. S.P., F.M.-N., A.L., L.D., N.L., L.G. performed statistical analyses. S.P., F.M.-N. and F.L. drafted the manuscript. All authors read and approved the final paper.

## Acknowledgements

The authors thank Cédric Van Marcke de Lummen (UCLouvain, Brussels, Belgium) for advice; Atsushi Kumanogoh (Osaka University, Osaka, Japan), Thomas Worzfeld (University of Marburg, Marburg, Germany) and Svetlana Chapoval, University of Maryland, Baltimore, MA, USA) for information on SEMA4A biology; the Lemaigre lab members for help and support.

