## Supplementary Material for "Spatial transcriptomics profiling of gallbladder adenocarcinoma: a detailed two-case study of progression from precursor lesions to cancer"

**SUPPLEMENTARY METHODS**

**Spatial transcriptomics.** Spatial profiling was performed on 5 μm-thick formalin-fixed paraffin-embedded (FFPE) tissue sections using GeoMx technology (NanoString Technologies, Seattle, WA, USA) [1] which was implemented by NanoString. FFPE human gallbladder slides were baked for 1 h at 65°C, and subsequently processed on Leica automation platform with a protocol that includes three steps: slide baking, antigen retrieval for 20 min at 100°C, and 1.0 μg/ml proteinase K treatment for 15 min. After taking the slides off the Leica automation platform, each slide was incubated overnight with the GeoMx Whole Transcriptome Atlas assay probe cocktail which contains 18,677 probes. The following day, the slides were washed and immunostained for markers of epithelium (anti-panCytokeratin), leukocytes (CD45), α smooth muscle actin (αSMA)-expressing mesenchymal cells, and cell nuclei (HuR), using the following antibodies: anti-CD45 (CST, #13917); anti-αSMA (AbCam, #ab202368); anti-HuR (Santa Cruz, #SC5261); anti-panCytokeratin (Novus, NBP2-33200AF488). Fluorescently-labeled slides were loaded onto a GeoMx machine and were scanned for selection of regions of interest (ROIs). Barcodes were collected and subsequent barcoding reading was performed on Illumina NGS platform.

**Immunofluorescence and immunohistochemistry.**

Supplementary Table S1. List of primary and secondary antibodies

| Primary antibody | Host | Company and reference | Dilution |
|---|---|---|---|
| CK7 | Mouse | Leica Biosystems #CK7-OVTL-L-CE | 1:100 |
| E-CADHERIN | Mouse IgG2 | BD #610182 | 1:300 |
| SM22$\alpha$ | Rabbit | Proteintech #10493-1-AP | 1:50 |
| SOX9 | Rabbit | Abcam #ab5535 | 1:100 |

| Secondary antibody | Host | Company and reference | Dilution |
|---|---|---|---|
| Alexa Fluor 488 anti-rat | Donkey | ThermoFisher Scientific #A-21208 | 1:500 |
| Alexa Fluor 488 mouse igG2a | Goat | ThermoFisher Scientific A-21131 | 1:500 |
| Alexa Fluor 594 anti-mouse | Donkey | ThermoFisher Scientific #A-21203 | 1:500 |
| Alexa Fluor 594 anti-rabbit | Donkey | ThermoFisher Scientific A-21207 | 1:500 |

**Gallbladder organoid culture.** Human non-tumoral gallbladder tissues were obtained from patients who underwent cholecystectomy at the Cliniques Universitaires Saint-Luc, Brussels, Belgium. We adapted the method of Rimland and coworkers [2]. Gallbladders were drained of bile and a small piece of the gallbladder body was excised, washed with Hanks Balanced Salt Solution (HBSS; Gibco/Thermo Fisher Scientific, Brussels, Belgium) and scraped along the luminal mucosal surface to mechanically dissociate the epithelium in Advanced DMEM/F-12 medium (Gibco). The cell suspension was washed twice through centrifugation at 300 g for 5 min and the cell pellet was resuspended in growth factor-reduced Matrigel (Corning/Merck, Hoeilaart, Belgium) before plating in droplets of 50 µl in a 24-well plate. After Matrigel solidification, media was added in the presence of 10 µM ROCK inhibitor Y-27632 (SigmaAldrich/Merck, Hoeilaart, Belgium) for the first 48 h after isolation or passage. The organoid culture medium consisted of 500 μl Advanced DMEM/F-12 containing 1x N2-serum free supplement (Gibco), 1x B27-serum free supplement (Gibco), 2 mM L-glutamine (Gibco), 1% Penicillin-Streptomycin (Gibco), 20% R-Spondin-conditioned media, 3 μM CHIR 99021 (Sigma), 100 ng/ml recombinant human noggin (SigmaAldrich), 2.5 μM PGE2 (R&D Systems/Bio-Techne, Abingdon, United Kingdom), 100 ng/ml recombinant human Epidermal Growth Factor (EGF; R&D Systems), 5 μM A 83-01 (PreproTech/Thermo Fisher Scientific, Brussels, Belgium) and 10 μM forskolin (MedChem Express/Bio-Connect, Huissen, The Netherlands). Media was changed every 2-3 days.

Four organoid lines were developed. The first line displayed trisomy 20 and 16q deletion, the second and third lines failed to expand. The fourth line grew as expected and was selected for further studies. Karytoping (GTG banding) of this line revealed no anomaly and was performed as described [3] with the following adaptations: Cells were centrifuged at 405 g, and slides were prepared in a temperature- and humidity-controlled environment chamber. HBSS was replaced with PBS, and a 2.5% trypsin solution (Thermo Fisher Scientific) was used. Slides were dried from 16 to 24 h in a dry oven at 70°C before staining with a 1.5% Giemsa solution (CellaVision - RAL Diagnostics, Martillac France). The Permount mounting medium was replaced with DPX (SigmaAldrich), and slides were briefly immersed in toluene (VWR Chemicals/Avantor, Leuven, Belgium) before the embedding step. Metaphases were selected with the Metafer scanning platform (v4.3; MetaSystems/Amplitech Cormeilles-en-Parisis, France) and analysed with Ikaros software (v6.3; MetaSystems). The coding-region of the genome of the organoid was analysed by whole exome sequencing (WES). Briefly, DNA was extracted using a DNeasy blood and tissue kit (ID 69504, Qiagen, Antwerp, Belgium). Quality controls and WES were performed by Macrogen Europe (Amsterdam, The Netherlands). Briefly, two (TWIST Core Exome plus RefSeq) libraries from the same organoid line were sequenced. Raw data (.fasta files) were aligned to the reference human genome assembly (GRCh38) using BWA 0.7.15 (Li and Durbin, Bioinformatics 2009), with the two libraries submitted together as different read groups. Aligned sequences (.bam file) were processed using Samtools 1.12 "MarkDup" (for marking duplicates) and GATK 4.2 "BQSR" (for base quality scores recalibration). This yielded a coverage depth of 332X (with duplicates removed). Germline single-nucleotide variants and small indels were identified using GATK 4.2 "Haplotype Caller" while somatic single-nucleotide variants (SNV) and small indels were identified using Mutect2 (both following Broad Institute best practices). The variant calls (.vcf) files generated were annotated, imported and further analyzed on Highlander 17.18 (https://sites.uclouvain.be/highlander/), the in-house bioinformatics framework of the Genomics and Bioinformatics platform of UCLouvain (PGEN: https://www.deduveinstitute.be/pgen-bioinformatics) for variant annotation, filtering and visualization, with a local database currently containing more than 4300 WES categorized by pathology. We filtered for variants in genes described as frequently mutated in gallbladder cancer according to Kuipers *et al.* [4]. An ERBB3$^{R675G}$ missense mutation was detected, at an allelic fraction of 0.021.

To analyse the impact of blocking SEMA4A in gallbladder organoids, the latter were split and plated. After 24 h, SEMA4A antibody (IgG-SEMA4A, #14-1002-82 eBioscience/Thermo Fisher scientific, Brussels, Belgium) was added into the medium (10 μg/ml) and organoids were grown for 3 days. Prior incubation, SEMA4A antibody was desalted using a Zeba Spin Desalting Column (Thermo Fisher Scientific).

**RNA extraction and Reverse Transcription-quantitative PCR.** Human gallbladder organoids were purified fom the Matrigel dome using Cell Recovery Solution (Corning) and RNA was extracted using RNAqueous™-Micro Total RNA Isolation Kit (Thermo Fisher Scientific) according to the manufacturer's protocol. cDNA synthesis was performed with MMLV reverse transcriptase (Thermo Fisher Scientific) according to manufacturer's protocol. Gene expression was quantified by qPCR using Kappa Syberfast qPCR kit (Sopachem, Nazareth, Belgium) using the following conditions: 3 min at 95°C, 40 cycles of denaturation (3 sec at 95°C), annealing and extension (30 sec at 60°C). *RPL19* was used as internal control.

Oligonucleotides used in qPCR are described in the table below. Quantification of gene expression was performed using the 2(-Delta Delta C(T)) method and was expressed as a ratio of target to *RPL19* levels. Each amplification was performed in duplicate.

| **Target** | | **Forward Primer** | **Reverse Primer** |
|---|---|---|---|
| RPL19 | H | 5'-CGAATGCCAGAGAAGGTCAC-3' | 5'-CCATGAGAATCCGCTTGTTT-3' |
| SEMA4A | H | 5'-AATGGCTCCCTCTTGCTGAT-3' | 5'-GACTGGAGGTGTTGCTCTCT-3' |
| PLXNB1 | H | 5'-CCTCACCTTTGATGGGACCT-3' | 5'-GCACATCACACCAGAACCAG-3' |

pan-cytokeratin / CD45 / aSMA / HuR

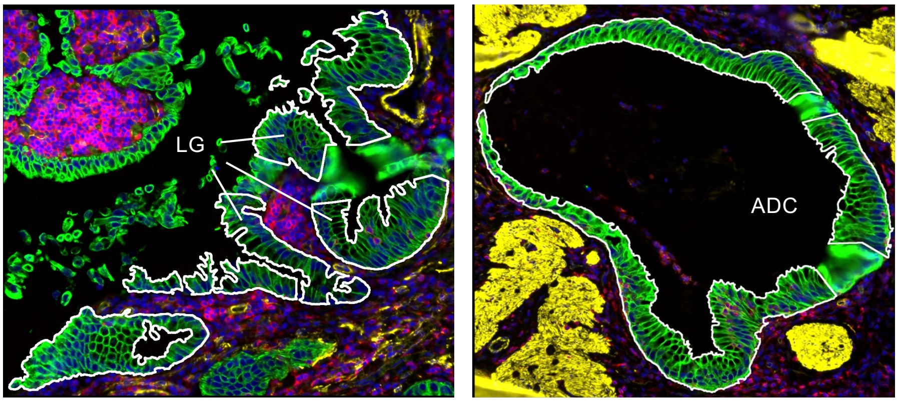


**Supplementary Fig. S1. Delineation of ROIs for spatial transcriptomic analysis.** Illustrative examples of gallbladder sections of Patient #2 with delineation (white line) of epithelial ROIs subjected to spatial transcriptomic analysis. ADC, area with adenocarcinoma; LG, area with low-grade biliary intraepithelial neoplasia (BilIN); NI, area with histologically normal epithelium.

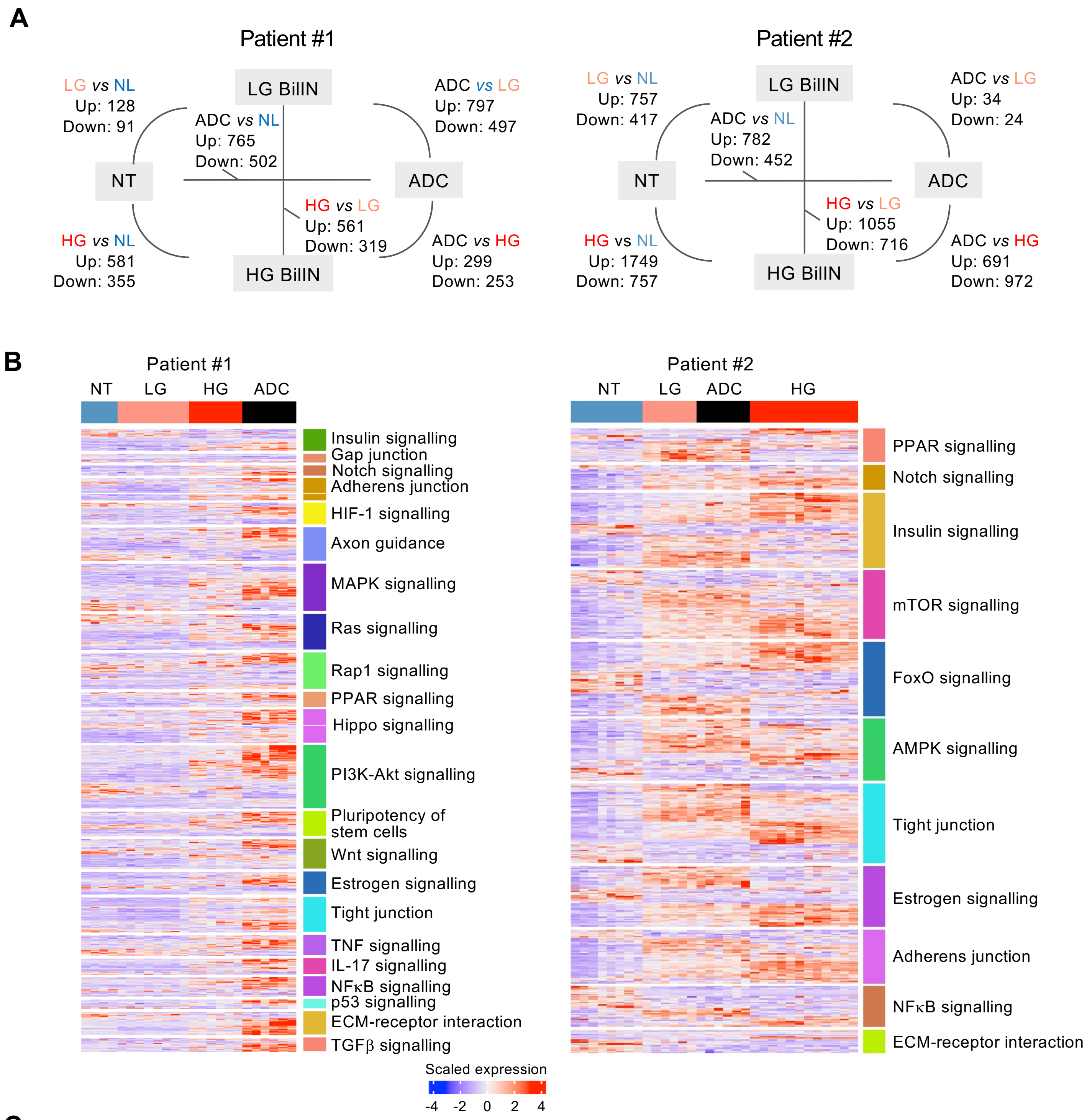


**C**

ADC *vs* Non-tumoral

Patient #1

| Gene | $\log_2$ fold | $p_{adj}$ |
|---|---|---|
| WNT7B | 0.85 | $2.52 \times 10^{-2}$ |
| WNT8A | 0.60 | $2.04 \times 10^{-2}$ |
| WNT10A | 1.25 | $3.54 \times 10^{-6}$ |
| WNT11 | 0.97 | $2.57 \times 10^{-2}$ |
| FZD2 | 1.35 | $8.05 \times 10^{-3}$ |
| FZD5 | 1.25 | $1.63 \times 10^{-3}$ |
| TCF7L2 | 1.14 | $7.12 \times 10^{-5}$ |
| AXIN2 | 0.82 | $3.90 \times 10^{-2}$ |
| GSK3B | 1.03 | $5.03 \times 10^{-4}$ |

**Supplementary Fig. S2: Differential gene expression in spatial transcriptomics. A** Number of genes up- or downregulated in cross-comparison of tissue types ($\log_2$ fold change ≥ 1.0; $p_{adj} \leq 0.05$). **B** Heatmap of genes from the KEGG pathway gene sets that are differentially expressed between adenocarcinoma and normal epithelium ROIs in the two patients ($p_{adj} \leq 0.05$). **C** List of upregulated Wnt signalling genes in Patient #1. ADC, adenocarcinoma; HG, high-grade biliary intraepithelial neoplasia (BilIN); LG, low-grade BilIN; NT, non-tumoral epithelium.

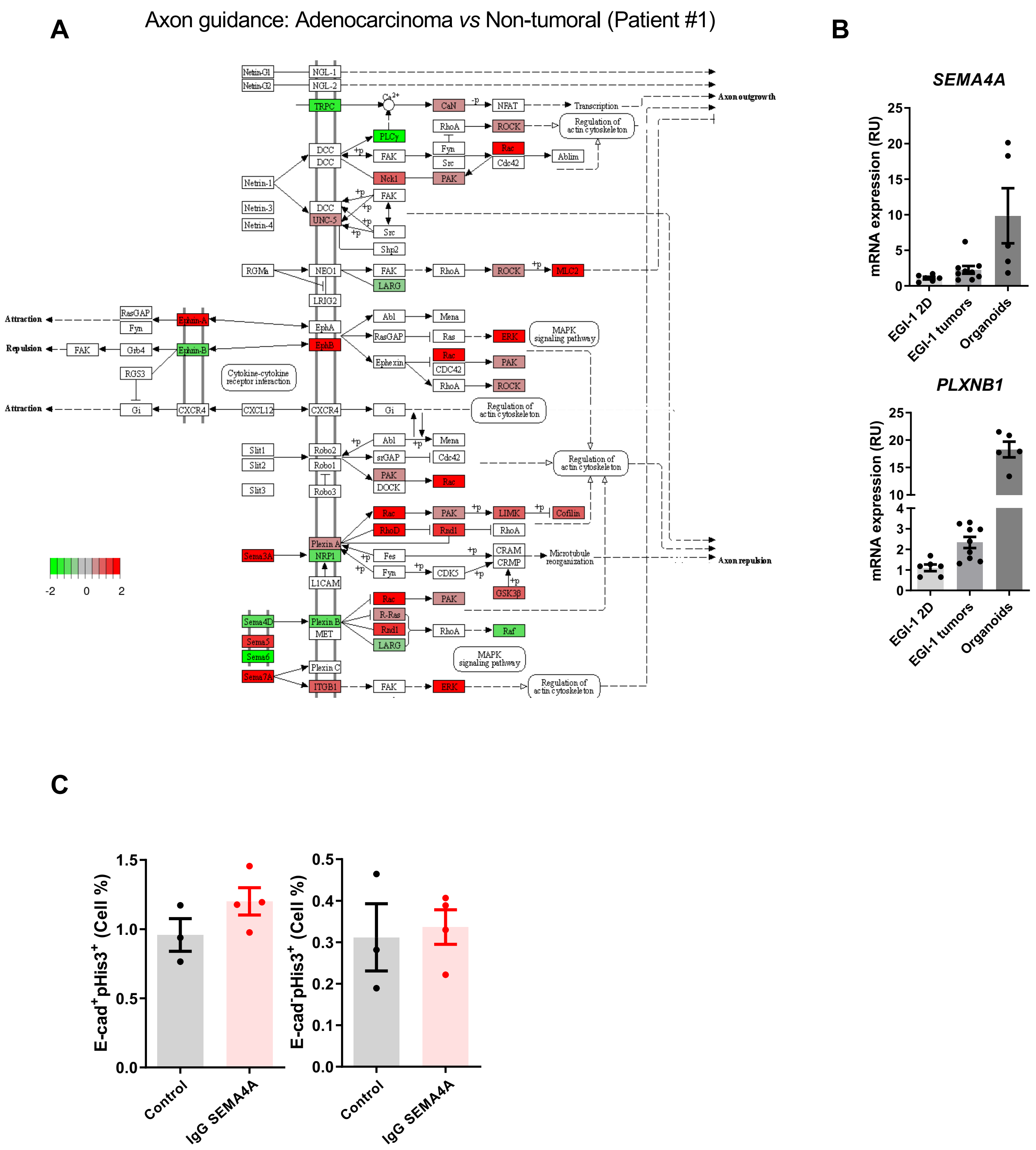


**Supplementary Fig. S3: Expression of axon guidance genes.** **A** Axon guidance pathway (KEGG) enriched in Patient #1 during tumour progression. **B** Expression of SEMAPHORIN 4A (SEMA4A) and its receptor PlexinB1 (PLXNB1) in cultured EGI-1 cells, EGI-1-derived subcutaneous tumours, and cultured galbladder organoids, as measured by RT-qPCR. SEMA4 and PLXNB1 primers are human-specific and do not detect potential expression in murine cells colonising EGI-1-cell derived tumours. **C** Percentage of E-CADHERIN (E-CAD)-positive cells and E-CADHERIN-negative cells expressing phospho-Histone H3 (pHis3) in EGI-1-cell-derived tumours at day 8 of treatment with PBS (= control) or blocking anti-SEMA4A IgG antibodies. NES, normalised enrichment score.

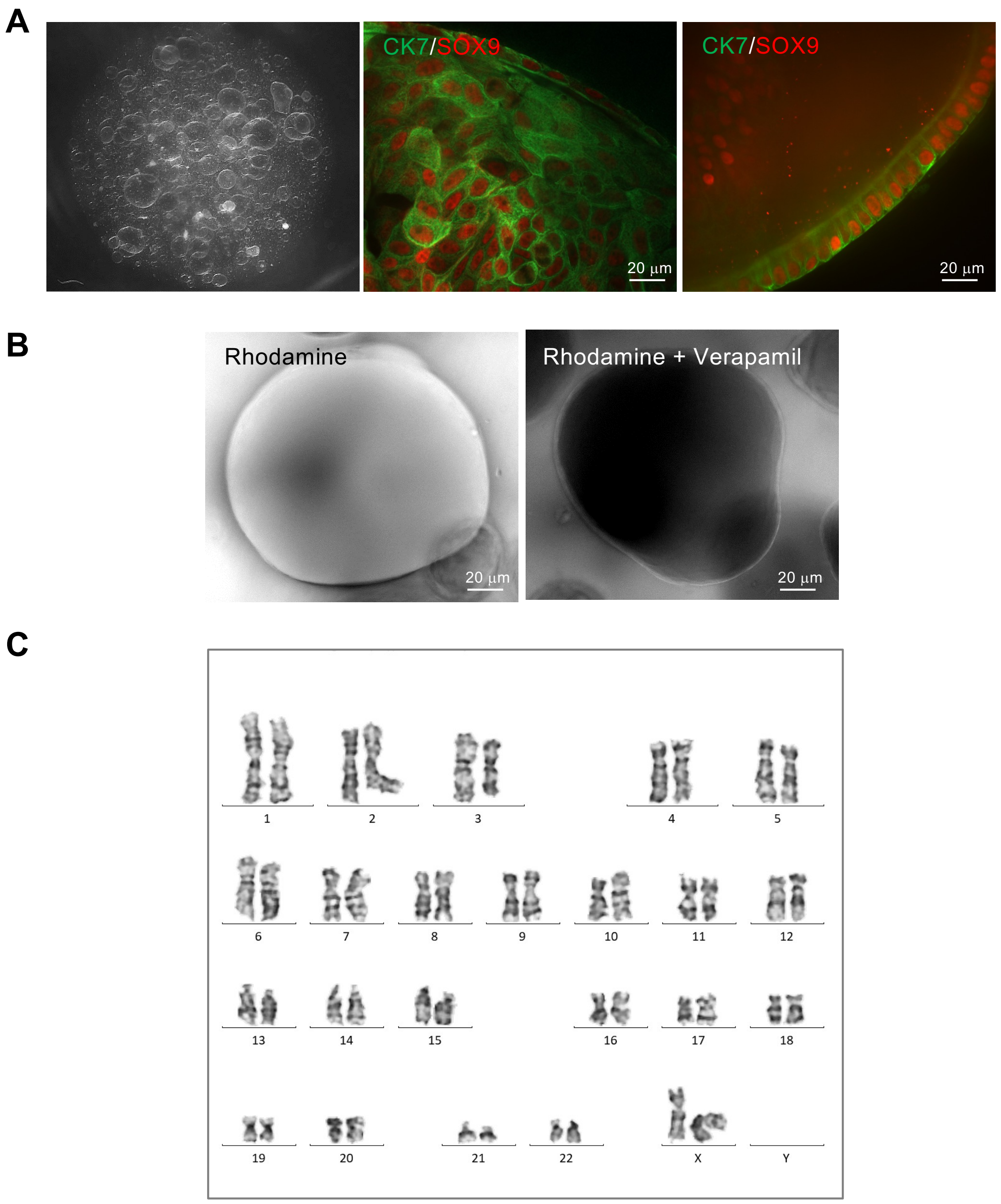


**Supplementary Fig. S4. Characterization of gallbladder organoids. A** Gallbladder organoids were grown in Matrigel and visualised using light microscopy (left panel). Immunostaining revealed expression of the biliary-specific proteins CYTOKERATIN 7 (CK7) and SRY-related HMG box transcription factor 9 (SOX9). **B** Intraluminal accumulation of Rhodamine (100 µM) that was added to the organoid medium was inhibited by Verapamil (10 µM), demonstrating the functionality of MDR1-mediated transport. **C** Normal karyotype of the selected organoid line.
